# Task demands affect spatial reference frame weighting during tactile localization in sighted and congenitally blind adults

**DOI:** 10.1101/056515

**Authors:** Jonathan T.W. Schubert, Stephanie Badde, Brigitte Röder, Tobias Heed

## Abstract

Task demands modulate tactile localization in sighted humans, presumably through weight adjustments in the spatial integration of anatomical, skin-based, and external, posture-based information. In contrast, previous studies have suggested that congenitally blind humans, by default, refrain from automatic spatial integration and localize touch using only skin-based information. Here, sighted and congenitally blind participants localized tactile targets on the palm or back of one hand, while ignoring simultaneous tactile distractors at congruent or incongruent locations on the other hand. We probed the interplay of anatomical and external location codes for spatial congruency effects by varying hand posture: the palms either both faced down, or one faced down and one up. In the latter posture, externally congruent target and distractor locations were anatomically incongruent and vice versa. Target locations had to be reported either anatomically (“palm” or “back” of the hand), or externally (“up” or “down” in space). Under anatomical instructions, performance was better for anatomically congruent than incongruent target-distractor pairs. In contrast, under external instructions, performance was better for externally congruent than incongruent pairs. These modulations were evident in sighted and blind individuals. Notably, distractor effects were overall far smaller in blind than in sighted participants, despite comparable target-distractor identification performance. Thus, the absence of developmental vision seems to be associated with an increased ability to focus tactile attention towards a non-spatially defined target. Nevertheless, that blind individuals exhibited effects of hand posture and task instructions in their congruency effects suggests that, like the sighted,, they automatically integrate anatomical and external information during tactile localization. Moreover, spatial integration in tactile processing is, thus, flexibly adapted by top-down information – here, task instruction – even in the absence of developmental vision.

## 1. Introduction

The brain continuously integrates information from multiple sensory channels (Alais & Burr, 2004; Angelaki, Gu, & DeAngelis, 2009; Ernst & Banks, 2002; Pouget, Deneve, & Duhamel, 2002; Sober & Sabes, 2005; Trommershäuser, Körding, & Landy, 2011). Tactile localization, too, involves the integration of several information sources, such as somatosensory information about the stimulus location on the skin with proprioceptive and visual information about the current body posture, and has therefore been investigated in the context of information integration within and across the senses. We have suggested that tactile localization involves at least two cortical processing steps (Badde, Heed, & Röder, 2015; Badde, Röder, & Heed, 2015). When tactile information first arrives in the cortex, it is initially encoded relative to the skin in an anatomical reference frame, reflected in the homuncular organization of the somatosensory cortex (Penfield & Boldrey, 1937). This information is consecutively remapped into an external reference frame. By merging anatomical skin-based spatial information with proprioceptive, visual, and vestibular signals, the brain derives an external spatial location, a process referred to as tactile remapping (Clemens, Vrijer, Selen, Gisbergen, & Medendorp, 2011; Driver & Spence, 1998; Heed, Buchholz, Engel, & Röder, 2015; Holmes & Spence, 2004; Maravita, Spence, & Driver, 2003). The term, external’, in this context, denotes a spatial code that abstracts from the original location, but is nevertheless egocentric, and takes into account the spatial relation of the stimulus to the eyes, head, and torso (Heed, Backhaus, Röder, & Badde, 2016; Heed, Buchholz, et al., 2015). In a second step, information coded with respect to the different reference frames is integrated, presumably to derive an superior tactile location estimate (Badde & Heed, 2016). For sighted individuals, this integration of different tactile codes appears mandatory (Azañón, Camacho, & Soto-Faraco, 2010; Shore, Spry, & Spence, 2002; Yamamoto & Kitazawa, 2001). Yet, the relative weight of each code is subject to change depending on current task demands: external spatial information is weighted more strongly when task instructions emphasize external spatial aspects (Badde & Heed, 2016; Badde, Heed, et al., 2015; Badde, Röder, et al., 2015), in the context of movement (Gherri & Forster, 2012a, 2012b; Heed, Möller, & Röder, 2015; Hermosillo, Ritterband-Rosenbaum, & van Donkelaar, 2011; Mueller & Fiehler, 2014a, 2014b; Pritchett, Carnevale, & Harris, 2012), and in the context of frequent posture changes (Azañón, Stenner, Cardini, & Haggard, 2015). Thus, the tactile localization estimate depends on flexibly weighted integration of spatial reference frames.

Moreover, tactile localization critically depends on visual input after birth. Differences between sighted and congenitally blind participants in touch localization are evident, for instance, in tasks involving hand crossing. Anatomical and external spatial reference frames can be experimentally misaligned by crossing the hands over the midline, so that the left hand occupies the right external space and vice versa. Hand crossing reportedly impairs tactile localization compared to an uncrossed posture in sighted, but not in congenitally blind individuals (Collignon, Charbonneau, Lassonde, & Lepore, 2009; Röder, Rösler, & Spence, 2004). Similarly, hand crossing can attenuate spatial attention effects on somatosensory event-related potentials (ERP) between approximately 100 and 250 ms post-stimulus in sighted, but not in congenitally blind individuals (Röder, Föcker, Hötting, & Spence, 2008). Together, these previous studies indicate that congenitally blind individuals may not integrate externally coded information with anatomical skin-based information by default when they process touch, setting them apart from sighted individuals. Recent studies, however, have cast doubt on this conclusion. For instance, congenitally blind individuals used external along with anatomical coding when tactile stimuli had to be localized while making bimanual movements (Heed, Möller, et al., 2015). Evidence for automatic integration of external spatial information in congenitally blind individuals comes not only from tactile localization, but, in addition, from a bimanual coordination task: when participants moved their fingers symmetrically, this symmetry was encoded relative to external space rather than according to anatomical parameters such as the involved muscles, similar as in sighted participants (Heed & Röder, 2014). In addition, it has recently been reported that early blind individuals encode locations of motor sequences relative to both external and anatomical locations (Crollen, Albouy, Lepore, & Collignon, 2013). Moreover, early blind individuals appear to encode time relative to external space, and this coding strategy may be related to left-right finger movements during Braille reading (Bottini, Crepaldi, Casasanto, Crollen, & Collignon, 2015). These studies imply that congenitally blind humans, too, integrate spatial information coded in different reference frames according to a weighting scheme (Badde & Heed, 2016), but use lower default weights for externally coded information than sighted individuals. In both groups, movement contexts seem to induce stronger weighting of external spatial information.

In sighted individuals, task demands are a second factor besides movement context that can modulate the weighting of spatial information in tactile localization. For instance, tactile temporal order judgments (TOJ), that is, the decision which of two tactile locations was stimulated first, are sensitive to the conflict between anatomical and external locations that arises when stimuli are applied to crossed hands. This crossing effect was modulated by a secondary task that accentuated anatomical versus external space, indicating that the two tactile codes were weighted according to the task context (Badde, Röder, et al., 2015). The modulatory effect of task demands on tactile spatial processing in sighted individuals is also evident in the tactile congruency task: In this task, tactile distractors presented to one hand interfere with elevation judgements about simultaneously presented tactile target stimuli presented at the other hand (Gallace, Soto-Faraco, Dalton, Kreukniet, & Spence, 2008; Soto-Faraco, Ronald, & Spence, 2004). In such tasks, one can define spatial congruency between target and distractor in two ways: In an anatomical reference frame two stimuli are congruent if they occur at corresponding skin locations; in an external reference frame two stimuli are congruent if they occur at corresponding elevations. If one places the two hands in the same orientation, for instance, with both palms facing down, anatomical and external congruency are in correspondence. However, when the palm of one hand faces up and the other down, two tactile stimuli presented at upper locations in external space will be located at incongruent anatomical skin locations, namely the palm of one, and back of the other hand. Thus, anatomical and external congruency do not correspond in this latter posture. A comparison of congruency effects between these postures provides a measure of the weighting of anatomical and external tactile codes. Whether congruency effects in this task were encoded relative to anatomical or relative to external space was modifiable by both task instructions and response modalities (Gallace et al., 2008). This modulation suggests that the weighting of anatomical and external spatial information in the tactile congruency task was flexible, and was modulated by task requirements.

With respect to congenitally blind humans, evidence as to whether task instructions modulate spatial integration in a similar way as in sighted individuals is currently indirect. Effects of task context have been suggested to affect reaching behavior of blind children (Millar, 1981). For tactile integration, one piece of evidence in favor of an influence of task instructions comes from the comparison of two very similar studies that have investigated tactile localization in early (Eardley & van Velzen, 2011) and congenitally blind humans (Röder et al., 2008) by comparing somatosensory ERPs elicited by tactile stimulation in different hand postures. Both studies asked participants to report infrequent tactile target stimuli on a pre-cued hand, but observed contradicting results: One study reported an attenuation of spatial attention-related somatosensory ERPs between 140 and 300 ms post-stimulus to non-target stimuli with crossed compared to uncrossed hands (Eardley & van Velzen, 2011), suggesting that external location had affected tactile spatial processing in early blind participants. The other study (Röder et al., 2008), in contrast, did not observe any significant modulation of spatial attention-related somatosensory ERPs by hand posture and concluded that congenitally blind humans do not, by default, use external spatial information for tactile localization. The two studies differed only in how participants were instructed about the to-be-monitored location. In the first study, the pitch of an auditory cue indicated the task-relevant side relative to external space in each trial (Eardley & van Velzen, 2011). In the second study, in contrast, the pitch of a cuing sound referred to the task-relevant hand, independent of hand location in external space (Röder et al., 2008). Thus, one may hypothesize that task instructions modulate how anatomical and external information is weighted in congenitally blind individuals as they do in the sighted.

Here, we investigated the weighting of anatomical and external reference frames by means of an adapted version of the tactile congruency task (Gallace et al., 2008; Soto-Faraco et al., 2004). Sighted and congenitally blind participants localized vibro-tactile target stimuli, presented randomly on the palm or back of one hand, while ignoring vibro-tactile distractors on the palm or back of the other hand. Thus, distractors could appear at an anatomically congruent or incongruent location. Hand posture was varied with either both palms facing downwards, or one palm facing downwards and the other upwards. With differently oriented hands, anatomically congruent stimuli were incongruent in external space and vice versa. We used this experimental manipulation to investigate the relative importance of anatomical and external spatial codes during tactile localization.

We introduced two experimental manipulations to investigate the role of task demands on the weighting of anatomical and external spatial information: a change of task instructions and a change of the movement context.

For the manipulation of task instructions, every participant performed two experimental sessions. In one session, responses were instructed anatomically, that is, with respect to palm or back of the hand. In a second session, responses were instructed externally, that is, with respect to upper and lower locations in space. We hypothesized that each task instruction would emphasize the weighting of the corresponding reference frame. This means that with differently oriented hands (that is, when anatomical and external reference frames are misaligned) the size, or even direction, of the congruency effect should depend on task instructions.

With the manipulation of movement context, we aimed at corroborating previous results suggesting that movement planning and execution as well as frequent posture changes lead to emphasized weighting of external spatial information (Azañón et al., 2015; Gherri & Forster, 2012a, 2012b; Heed, Möller, et al., 2015; Hermosillo et al., 2011; Mueller & Fiehler, 2014a, 2014b; Pritchett et al., 2012). Accordingly, we hypothesized that frequent posture changes would increase the weight of the external reference frame in a similar way for the spatial coding of congruency in the present task. To this end, participants either held their hands in a fixed posture for an entire experimental block, or they changed their hand posture in a trial-by-trial fashion. Again, with differently oriented hands, changes in the weighting of anatomical and external spatial information would be evident in a modulation of tactile congruency effects. If frequent posture changes, compared to a blockwise posture change, induce an increased weighting of external information, this would result in a decrease of anatomical congruency effects under anatomical instructions and in an increase of external congruency effects under external instructions.

## 2. METHODS

We follow open science policies as suggested by the Open Science Framework (see https://osf.io/hadz3/wiki/home/) and report how we determined the sample size, all experimental manipulations, all exclusions of data, and all evaluated measures of the study. Data and analysis scripts are available online (see https://osf.io/ykqhd/). For readability, we focus on accuracy in the paper, but report reaction times and their statistical analysis in the supporting information. Previous studies have reported qualitatively similar results for both measures, and results were also comparable in the present study.

### 2.1. Participants

The size of our experimental groups was constrained by the availability of congenitally blind volunteers; we invited every suitable participant we identified within a period of 6 months. Group size is comparable to that of previous studies that have investigated spatial coding in the context of tactile congruency. We report data from sixteen congenitally blind participants (8 female, 15 right handed, 1 ambidextrous, age: M = 37 years, SD = 11.6, range: 19 to 53) and from a matched control group of sixteen blindfolded sighted participants (8 female, all right handed, age: M = 36 years, SD = 11.5, range: 19 to 51). All sighted participants had normal or corrected-to-normal vision. Blind participants were visually deprived from birth due to anomalies in peripheral structures resulting either in total congenital blindness (n = 6) or in minimal residual light perception (n = 10). Peripheral defects included binocular anophthalmia (n = 1), retinopathy of prematurity (n = 4), Leber’s congenital amaurosis (n = 1), congenital optical nerve atrophy (n = 2), and genetic defects that were not further specified (n = 8). All participants gave informed written consent and received course credit or monetary compensation for their participation. The study was approved by the ethical board of the German Psychological Society (TB 122010) and conducted in accordance with the standards laid down in the Declaration of Helsinki.

Of twenty originally tested congenitally blind participants, one did not complete the experiment, and data from three participants were excluded due to performance at chance level (see below). We recruited 45 sighted participants to establish a group of 16 control participants. Technical failure during data acquisition prevented the use of data from two out of these 45 participants. Furthermore, we had developed and tested the task at the beginning in a young, sighted student population. We then tested the blind participants before recruiting matched controls from the population of Hamburg via online and newspaper advertisement. For many sighted, age-matched participants older than 30 years, reacting to the target in the presence of an incongruent distractor stimulus in the tactile congruency task proved too difficult, resulting in localization performance near chance level. Accordingly, 23 sighted participants either decided to quit, or were not invited for the second experimental session because their performance in the first session was not sufficient. We address this surprising difference between blind and sighted participants in ability to perform the experimental task in the Discussion.

### 2.2. Apparatus

Participants sat in a chair with a small lap table on their legs. They placed their hands in a parallel posture in front of them, with either both palms facing down (termed “same orientation”) or with one hand flipped palm down and the other palm up (termed “different orientation”). Whether the left or the right hand was flipped in the other orientation condition was counterbalanced across participants. Distance between index fingers of the hands was approximately 20 cm measured while holding both palms down. For reasons of comfort and to avoid stimulators touching the table small foam cubes supported the hands. Custom-built vibro-tactile stimulators were attached to the back and to the palm of both hands midway between the root of the little finger and the wrist (Fig. 1). Participants wore earplugs and heard white noise via headphones to mask any sound produced by the stimulators. Hand posture was monitored using a movement tracking system (Visualeyez II VZ4000v PTI; Phoenix Technologies Incorporated, Burnaby, Canada), with LED markers attached to the palm and back of the hands. The experiment was controlled using Presentation (version 16.2; Neurobehavioral Systems, Albany, CA, USA), which interfaced with Matlab (The Mathworks, Natick, MA, USA) and tracker control software VZSoft (PhoeniX Technologies Incorporated, Burnaby, Canada).

**Figure 1.**
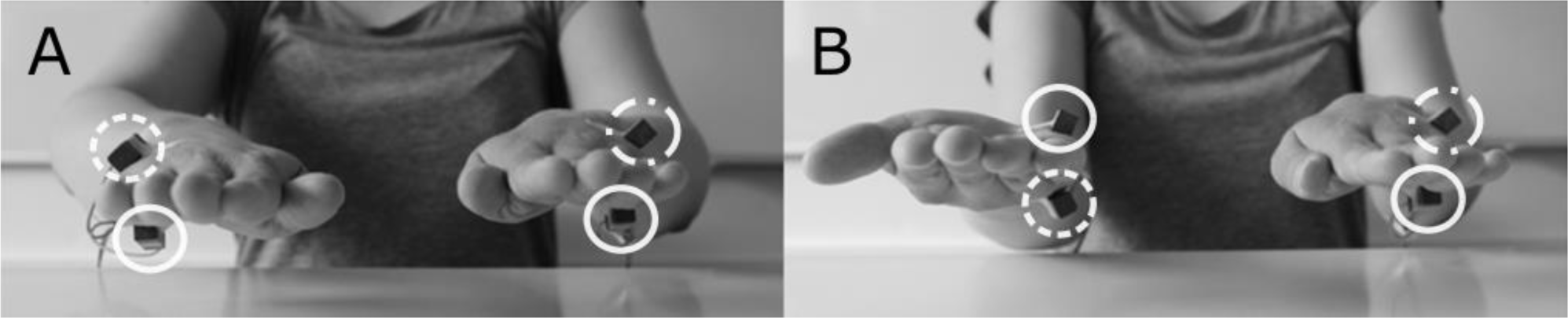
Experimental setup. Four vibro-tactile stimulators were attached to the palm and back of the hands (marked with white circles). The hands were either held in the same orientation with both palms facing downwards (**A**) or in different orientations with one hand flipped upside-down (**B**). In each trial, a target stimulus was randomly presented at one of the four locations. Simultaneously, a distractor stimulus was presented randomly at one of the two stimulator locations on the other hand. Target and distractor stimuli differed with respect to their vibration pattern. Participants were asked to localize the target stimulus as quickly and accurately as possible. For statistical analysis and figures, stimulus pairs presented to the same anatomical locations were defined as congruent, as illustrated by dashed (target) and dashed-dotted (distractor) circles, which both point to the back of the hand here. Note that with differently oriented hands (**B**) anatomically congruent locations are incongruent in external space and vice versa.

### 2.3. Stimuli

The experiment comprised two kinds of tactile stimuli, namely, targets, to which participants had to respond, and distractors, which participants had to ignore. Target stimuli consisted of 200 Hz stimulation for 235 ms. Distractor stimuli vibrated with the same frequency, but included two gaps of 65 ms, resulting in three short bursts of 35ms each. We initially suspected that our sighted control participants’ low performance in the congruency task may be related to difficulty in discriminating target and distractor stimuli. Therefore, we adjusted the distractor stimulus pattern for the last seven recruited control participants if they could not perform localization of the target above chance level in the presence of an incongruent distractor stimulus during a pre-experimental screening; such adjustments were, however, necessary for only three of these last seven participants, for the four other participants no adjustments were made. In a first step, we increased the distractor’s gap length to 75 ms, resulting in shorter bursts of 25 ms (1 participant). If the participant still performed at chance level in incongruent trials of the localization task, we set the distractor pattern to 50 ms “on”, 100 ms “off”, 5 ms “on”, 45 ms “off”, and 35 ms “on” (2 participants). Note that, while these distractor stimulus adjustments made discrimination between target and distractor easier, they did not affect target localization per se. Consequently, it is possible that localization performance in incongruent trials simply improved due to the additional training during stimulus adjustment. Importantly, however, stimuli were the same in all experimental conditions. Yet, to ascertain that statistical results were not driven by these three control participants, we ran all analyses both with and without their data. The overall result pattern was unaffected, and we thus report results of the full control group. In addition, we ascertained that sighted and blind groups could both discriminate target and distractor stimuli equally well. Practice before the task included training in stimulus discrimination, and discrimination performance during practice did not differ statistically between the two groups (see supporting information). Moreover, discrimination performance during practice and the size of congruency effects did not significantly correlate within each group (see supporting information).

### 2.4. Procedure

The experiment was divided into four large parts according to the combination of the two experimental factors Instruction (anatomical, external) and Movement Context (static vs. dynamic context, that is, blockwise vs. frequent posture changes). The order of these four conditions was counterbalanced across participants. Participants completed both Movement Context conditions under the first instruction within one session, and under the second instruction in a another session, which took part on another day. Participants completed four blocks of 48 trials for each combination of Instruction and Movement Context. Trials in which participants responded too fast (RT < 100 ms), or not at all, were repeated at the end of the block.

### 2.5. Manipulation of instruction

Under external instructions, participants had to report whether the target stimulus was located “up” or “down” in external space and ignore the distractor stimulus. They had to respond as fast and accurately as possible by means of a foot pedal placed underneath one foot (left and right counterbalanced across participants), with “up” responses assigned to lifting of the toes, and “down” responses to lifting of the heel. Previous research in our lab showed that participants strongly prefer this response assignment and we did not use the reverse assignment to prevent increased task difficulty. Note, that congruent and incongruent target-distractor combinations required up and down responses with equal probability, so that the response assignment does not bias our results. Under anatomical instructions, participants responded whether the stimulus had occurred to the palm or back of the hand. We did not expect a preferred response mapping under these instructions and, therefore, balanced the response mapping of anatomical stimulus location to toe or heel across participants.

### 2.6. Manipulation of movement context

Under each set of instructions, participants performed the entire task once with a constant hand posture during entire experimental blocks (static movement context), and once with hand posture varying from trial to trial (dynamic movement context).

#### 2.6.1. Static movement context

In the static context, posture was instructed verbally at the beginning of each block. A tone (1000 Hz sine, 100 ms) presented via loudspeakers placed approximately 1 m behind the participants signaled the beginning of a trial. After 1520 - 1700 ms (uniform distribution) a tactile target stimulus was presented randomly at one of the four locations. Simultaneously, a tactile distractor stimulus was presented at one of the two locations on the other hand. Hand posture was changed after completion of the second of four blocks.

#### 2.6.2. Dynamic movement context

In the dynamic context, an auditory cue at the beginning of each trial instructed participants either to retain (one beep, 1000 Hz sine, 100 ms) or to change (two beeps, 900 Hz sine, 100 ms each) the posture of the left or right hand (constant hand throughout the experiment, but counterbalanced across participants). After this onset cue, the trial continued only when the corresponding motion tracking markers attached to the hand surfaces had been continuously visible from above for 500 ms. If markers were not visible 5000 ms after cue onset, the trial was aborted and repeated at the end of the block. An error sound reminded the participant to adopt the correct posture. Tactile targets occurred equally often at each hand, so that targets and distracters, respectively, occurred half of the time on the moved, and half of the time on the unmoved, hand. The order of trials in which posture changed and trials in which posture remained unchanged, was pseudo-randomized in a way to assure equal amounts of trials for both conditions.

#### 2.7. Practice

Before data acquisition, participants familiarized themselves with the stimuli by completing two blocks of 24 trials in which each trial randomly contained either a target or a distractor, and participants reported with the footpedal which of the two had been presented. Next, participants localized 24 target stimuli without the presence of a distractor stimulus to practice the current stimulus-response mapping (anatomical instructions: palm and back of the hand vs. external instructions: upper or lower position in space to toes and heel). Finally, participants practiced five blocks of 18 regular trials, two with the hands in the same orientation, and three with the hands in different orientations. Auditory feedback was provided following incorrect responses during practice, but not during the subsequent experiment.

#### 2.8. Data analysis

Data were analyzed and visualized in R (version 3.2.2; R Core Team, 2015) using the R packages lme4 (v1.1-9; Bates, Maechler, Bolker, & Walker, 2014), afex (v0.14.2; Singmann, Bolker, & Westfall, 2015), lsmeans (v2.20-2; Lenth & Hervé, 2015), dplyr (v0.4.3; Wickham & Francois, 2015), and ggplot2 (v1.0.1; Wickham, 2009). Trials with reaction times longer than 2000 ms were excluded from further analysis (5.58% of all trials).

We analyzed accuracy using generalized linear mixed models (GLMM) with a binomial link function (Jaeger, 2008; Bolker et al., 2009). It has been suggested that a full random effects structure should be used for significance testing in (G)LMM (Barr, Levy, Scheepers, & Tily, 2013). However, it has been shown that conclusions about fixed effect predictors do not diverge between models with maximal and models with parsimonious random effects structure (Bates, Kliegl, Vasishth, & Baayen, 2015), and it is common practice to include the maximum possible number of random factors when a full structure does not result in model convergence (e.g., Bolker et al., 2009; Brandes & Heed, 2015). In the present study, models reliably converged when we included random intercepts and slopes per participant for each main effect, but not for interactions. Significance of fixed effects was assessed with likelihood ratio tests comparing the model with the maximal fixed effects structure and a model that excluded the fixed effect of interest (Pinheiro & Bates, 2000). These comparisons were calculated using the afex package (Singmann et al., 2015), and employed Type III sums of squares and sum-to-zero contrasts. Fixed effects were considered significant at p < 0.05. Post-hoc comparisons of significant interactions were conducted using approximate z-tests on the estimated least square means (LSM, lsmeans package; Lenth & Hervé, 2015). The resulting p-values were corrected for multiple comparisons following the procedure proposed by Holm (1979). To assess whether the overall result pattern differed between groups, we fitted a GLMM with the fixed between-subject factor Group (sighted, blind) and fixed within-subjects factors Instruction (anatomic, external), Posture (same, different), Congruency (congruent, incongruent), and Movement Context (static, dynamic). Congruency was defined relative to anatomical locations for statistical analysis and figures. Subsequently, to reduce GLMM complexity and to ease interpretability, we conducted separate analyses for each group including the same within-subject fixed effects.

### 3. RESULTS

We assessed how task instructions and movement context modulate the weighting of anatomically and externally coded spatial information in a tactile-spatial congruency task performed by sighted and congenitally blind individuals.

We report here on accuracy, but, in addition, present reaction times and their statistical analysis in the supporting information. Overall, reaction times yielded a result pattern comparable to that of accuracy. Weight changes should become evident in a modulation of congruency effects within hand postures that induce misalignment between these different reference frames. With differently oriented hands, stimulus pairs presented to anatomically congruent locations are incongruent in external space and vice versa, whereas the two coding schemes agree when the hands are in the same orientation. Thus, a modulation of reference frame weighting by task instructions would be evident in an interaction of Instruction, Posture, and Congruency. Furthermore, a modulation of weights by the movement context would be evident in an interaction of Movement Context, Posture, and Congruency.

Figure 2 depicts mean accuracy of sighted and congenitally blind groups. A GLMM (Table 1) with fixed effect factors Group, Instruction, Posture, Congruency, and Movement Context revealed a four-way interaction of Group, Instruction, Posture, and Congruency (χ^2^(1) = 13.83, p < 0.001) Successively, we analyzed accuracy separately for each group.

**Figure 2.**
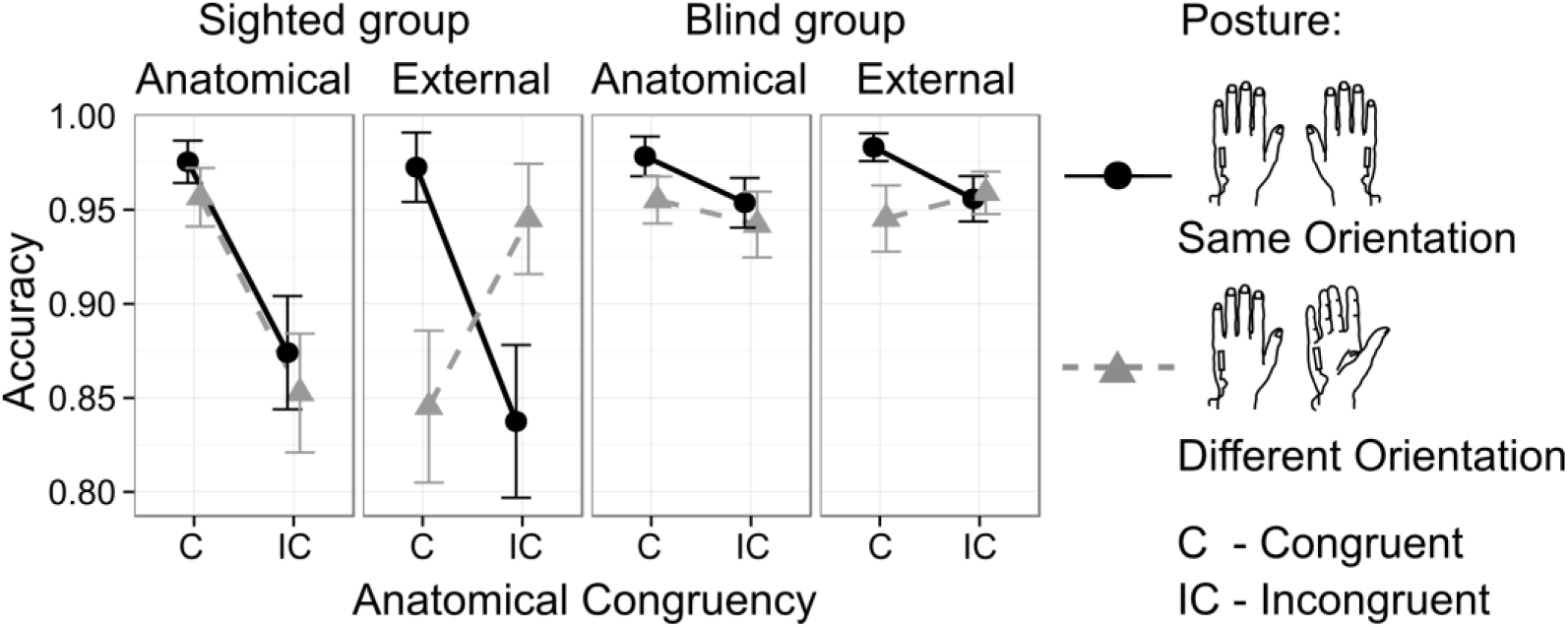
Accuracy in the tactile congruency task collapsed over static and dynamic movement conditions. Sighted (1^st^ and 2^nd^ column) and congenitally blind participants (3^rd^ and 4^th^ column) were instructed to localize tactile targets either relative to their anatomical (1^st^ and 3^rd^ column) or relative to their external spatial location (2^nd^ and 4^th^ column). Hands were placed in the same (black circles) and in different orientations (grey triangles). Tactile distractors were presented to anatomically congruent (C) and incongruent (IC) locations of the other hand and had to be ignored. Congruency is defined in anatomical terms (see Fig. 1). Accordingly, with differently oriented hands, anatomically congruent stimulus pairs are incongruent in external space and vice versa. Whiskers represent the standard error of the mean. Although accuracy was analyzed with a log-link GLMM, we present untransformed percentage-correct values to allow a comparison to previous studies (see methods for details).

**Table 1.**
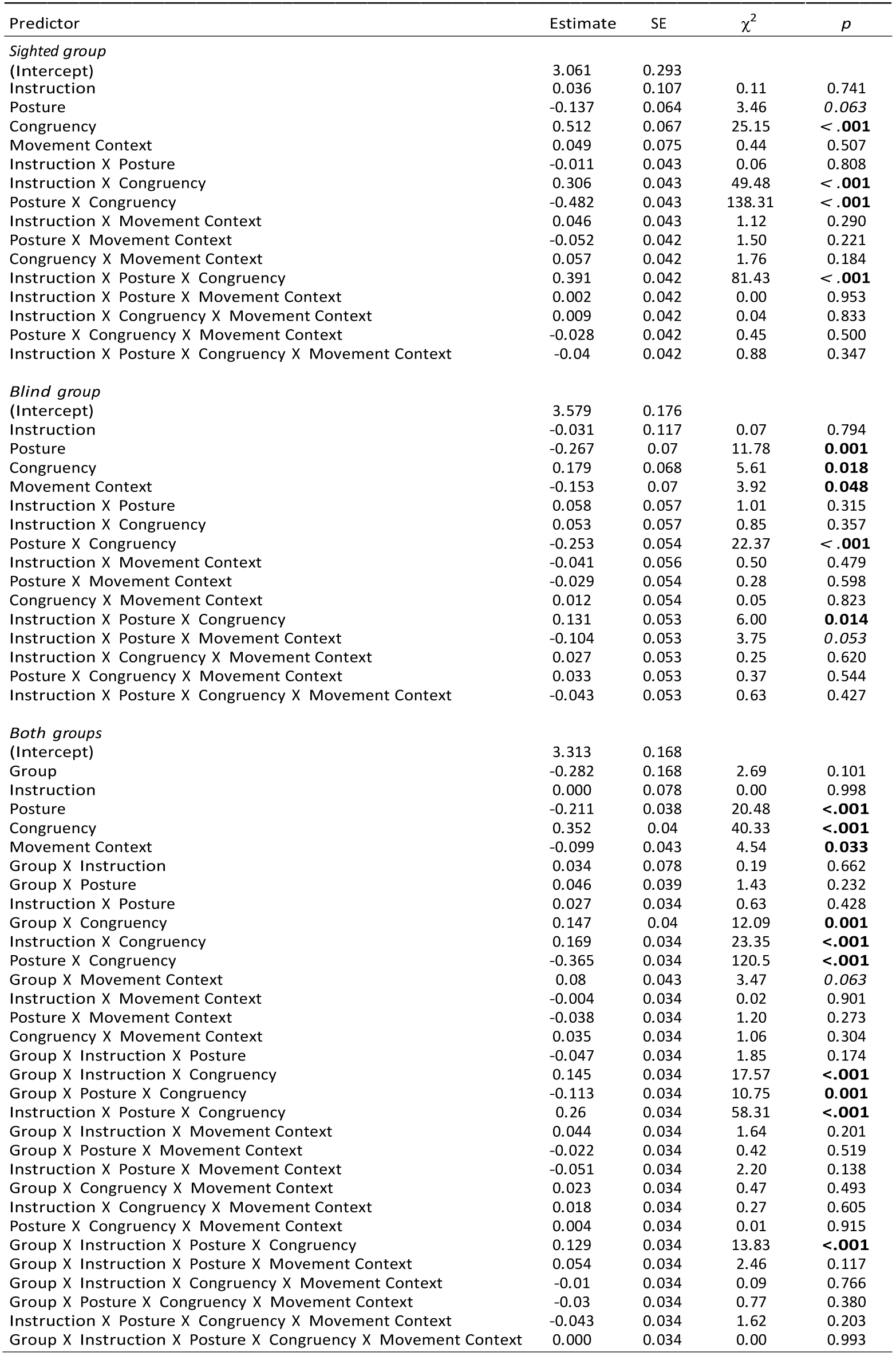
Statistical results from accuracy analysis. Summary of the fixed effects in the GLMM of the sighted group, of the blind group, and of the combined analysis. Coefficients are logit units. Bold values indicate significance at p < 0.05. Italic values indicate a trend towards significance at p < 0.1. Test statistics are χ^2^-distributed with 1 degree of freedom.

### 3.1. Sighted group: Manipulation of task instruction

The GLMM for the sighted group (Table 1) revealed a three-way interaction between Instruction, Posture, and Congruency (χ^2^(1) = 81.43, p < 0.001), suggesting that congruency effects differed in dependence of Instruction and Posture. Indeed, post-hoc comparisons revealed a two-way interaction between Posture and Congruency under external (z = 14.80, p < 0.001), but not under anatomical instructions (z = 1.50, p = 0.133). Under anatomical instructions, an anatomical congruency effect emerged across postures (Fig. 2, 1^st^ column; z = -10.15, p < 0.001). Under external instructions, participants responded more accurately following (anatomically and externally) congruent than incongruent stimulation when hands were oriented in the same orientation (Fig. 2, 2^nd^ column, black circles z = 10.26, p < 0.001). This congruency effect was reversed when the hands were held in different orientations, with more accurate performance for externally congruent (i.e., anatomically incongruent) than externally incongruent (i.e., anatomically congruent) stimulus pairs (Fig. 2, 2^nd^ column, gray triangles; z = -7.32, p < 0.001). Thus, the direction of the tactile congruency effect depended on the instructions. In sum, congruency effects were consistent with the instructed spatial coding – anatomical or external – in sighted participants.

### 3.2. Sighted group: Manipulation of movement context

Neither the effect of Movement Context (Fig. 3 top) nor interactions of Movement Context with any other variable were significant in the GLMM on accuracy (all χ^2^(1) ≤ 1.50, p ≥ 0.221). To demonstrate that these null effects were not due simply to high variance or a few outliers, Fig. 3 shows individual participants’ performance. In sum, movement context did not significantly affect the congruency effect.

**Figure 3.**
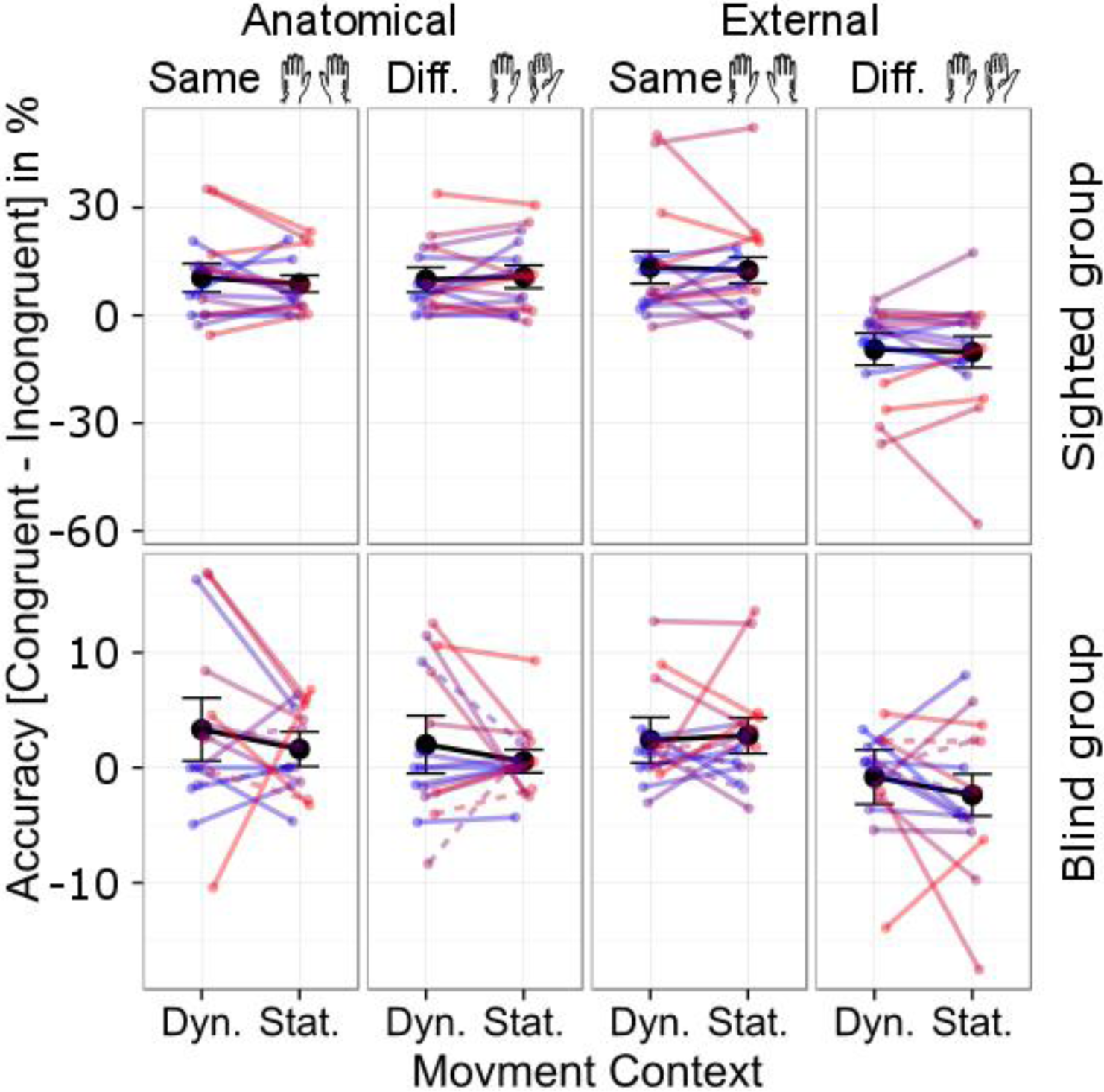
Individual participants’ tactile congruency effects in accuracy. Responses from anatomically incongruent trials were subtracted from responses in congruent trials. Congruency effects are plotted for dynamic (“Dyn.”) and static (“Stat.”) contexts with hands held in the same (1^st^ and 3^rd^ column) and in different orientations (2^nd^ and 4^th^ column) under anatomical (1^st^ and 2^nd^ column) and external instructions (3^rd^ and 4^th^ column) in the sighted (top row), and in the congenitally blind group (bottom row). Note that scales differ between groups, reflecting the smaller congruency effects of the blind as compared to the sighted group. Mean congruency effects for each condition are plotted in black, whiskers represent SEM. Each color represents one participant.

### 3.3. ongenitally blind group: Manipulation of task instruction

The GLMM on blind participants’ accuracy revealed a significant three-way interaction between Instruction, Posture, and Congruency (χ^2^(1) = 6.00, p = 0.014), suggesting that task instructions modulated the congruency effect. Post-hoc comparisons revealed a two-way interaction between Posture and Congruency under external instructions (z = 4.91, p < 0.001) and a trend for a two-way interaction under anatomical instructions (z = 1.66, p = 0.097). Under anatomical instructions, an anatomical congruency effect emerged across postures (Fig. 2, 3^rd^ column; z = -2.70, p = 0.007), and a main effect of Posture emerged across congruency conditions (z = 2.38, p = 0.017). Under external instructions, participants responded more accurately following (anatomically and externally) congruent than incongruent stimulation when hands were oriented in the same orientation (Fig. 2, 4^th^ column, black circles z = 3.75, p = 0.001). This congruency effect was reversed when the hands were held in different orientations, with more accurate performance for externally congruent (i.e., anatomically incongruent) than externally incongruent (i.e., anatomically congruent) stimulus pairs (Fig. 2, 4^th^ column, gray triangles; z = -2.56, p = 0.021). In sum, both task instruction and hand posture modulated congruency effects on accuracy in congenitally blind participants in a similar fashion as in sighted participants.

### 3.4. Congenitally blind group: Manipulation of movement context

The GLMM on blind participants’ accuracy showed a main effect of Movement Context (Fig.4, bottom; χ^2^(1) = 3.92, p = 0.048), with more accurate responses in the static than in the dynamic context. Moreover, there was a trend for a three-way interaction between Instruction, Posture, and Movement Context (χ^2^(1) = 3.75, p = 0.053; we report a follow-up analysis on this marginally significant trend in the supporting information). The GLMM did not yield any significant interactions involving Movement Context and Congruency. Thus, our hypothesis that a dynamic context modulates spatial integration in tactile congruency coding of congenitally blind humans did not receive any substantial support.

### 3.5. Comparison of the congruency effect between sighted and congenitally blind participants

Visual inspection of Figure 2 suggests that congruency effects were overall larger in the sighted than in the blind group; this result pattern was evident in reaction times as well (see supporting information). This difference in the size of congruency effects seemed to stem from the blind participants outperforming sighted participants when responding to stimulus pairs that were incongruent relative to the instructed reference frame (i.e., anatomically incongruent under anatomical instructions and externally incongruent under external instructions). To confirm this interaction effect statistically, we accordingly followed up on the significant four-way interaction between Group, Instruction, Posture, and Congruency (see Table 1) with post-hoc comparisons for each combination of Instruction and Posture. Indeed, there were significant two-way interactions between Group and Congruency, with larger congruency effects in the sighted than in the blind group for almost all combinations of Instruction and Posture (Table 2A, bold font). Moreover, blind participants responded more accurately than sighted participants following stimulus pairs that were anatomically incongruent under anatomical instructions and externally incongruent under external instructions (all p ≤ 0.016, bold font in Table 2B). In sum, congenitally blind participants exhibited smaller congruency effects than sighted participants due to better performance in incongruent conditions.

**Table 2.**
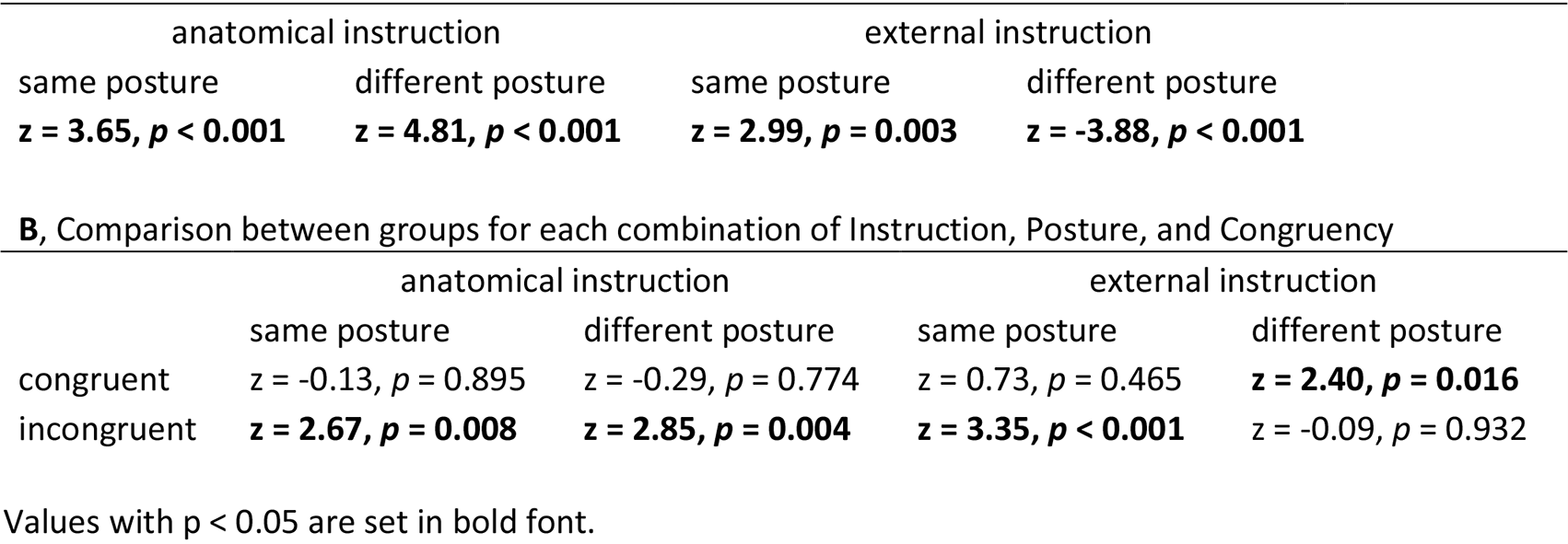
**A**, Interaction between Group and Congruency for each instruction and posture

## 4. DISCUSSION

The present study investigated whether congenitally blind individuals integrate anatomical and external spatial information during tactile localization in a flexible manner, similar to sighted individuals. To this end, participants localized tactile target stimuli at one hand in the presence of spatially congruent or incongruent distractor stimuli at the other hand. By manipulating hand posture, we varied the congruency of target and distractor locations relative to anatomical and external spatial reference frames. The study comprised two contextual manipulations, both of which influence tactile localization performance in sighted humans according to previous reports. First, we manipulated task context by formulating task instructions with reference to anatomical vs. external spatial terms (hand surfaces vs. elevation in space). Second, we manipulated movement context by maintaining participants’ hand posture for entire experimental blocks, or alternatively changing hand posture in a trial-by-trial fashion. Task instructions determined the direction of the congruency effect in both groups: Under anatomical instructions, both sighted and congenitally blind participants responded more accurately to anatomically congruent than incongruent target-distractor pairs, whereas under external instructions, participants responded more accurately to externally congruent than to externally incongruent target-distractor pairs. Even though blind participants exhibited congruency effects, interference from incongruent distractors was small compared to the sighted group due to superior performance in incongruent conditions. Movement context, that is, static hand posture versus frequent posture change, did not significantly modulate congruency effects in either experimental group.

### 4.1. Flexible weighting of reference frames according to task instructions in both sighted and blind individuals

Tactile externally coded spatial information has long been presumed to be automatically integrated only by normally sighted and late blind, but not by congenitally blind individuals, (Collignon et al., 2009; Röder et al., 2008, 2004). In the present study, blind participants’ performance should have been independent of posture and instructions if they had relied on anatomical information alone. In contrast to this assumption, blind participants’ congruency effects changed with posture when the task had been instructed externally, clearly reflecting the use of externally coded information.

The flexible and strategic weighting of anatomical and external tactile information, observed here in both sighted and blind individuals, presumably reflects top-down regulation of spatial integration (Badde & Heed, 2016). In line with this proposal, anatomical and external spatial information are presumed to be available concurrently, as evident, for instance, in event-related potentials (Heed & Röder, 2010) and in oscillatory brain activity (Buchholz, Jensen, & Medendorp, 2011, 2013; Schubert et al., 2015). Furthermore, performance under reference frame conflict, for instance due to hand crossing, is modulated by a secondary task, and this modulation reflects stronger weighting of external information when the secondary task accentuates an external as compared to anatomical spatial code (Badde, Röder, et al., 2015). The present results, too, demonstrate directed, top-down mediated modulations of spatial weighting, with anatomical task instructions biasing weighting towards anatomical coding, and external instructions biasing weighting towards external coding.

The present study’s results seemingly contrast with findings from previous studies. In several previous experiments, hand posture modulated performance of sighted, but not of congenitally blind participants. These results have been interpreted as indicating that sighted, but not congenitally blind individuals integrate external spatial information for tactile localization as a default (Collignon et al., 2009; Röder et al., 2008, 2004). In our experiment, both groups used external coding under external instructions and anatomical coding under anatomical instructions, indicating that developmental vision is not a prerequisite for the ability to flexibly weight information coded in the different reference frames. Thus, sighted and blind individuals do not seem to differ in their general ability to weight information from anatomical and external reference frames; instead, they appear to differ with respect to the reference frame they prefer under ambiguous task requirements. In other words, developmental vision seems to bias the default weighting of anatomical and external information towards the external reference frame, rather than changing the ability to adapt these weights.

### 4.2. Reduced distractor interference in congenitally blind individuals

Congenitally blind participants performed more accurately compared to sighted participants in the presence of incongruent distractors, both when target and distractor were anatomically incongruent under anatomical instructions, and when they were externally incongruent under external instructions. In addition, we had to exclude many sighted participants from the study because they performed at chance level when target and distractor were incongruent relative to the current instruction. Critically, this difficulty of sighted participants with the task cannot be due to an inability to discriminate between target and distractor stimuli: neither did discrimination performance significantly differ between sighted and blind groups, nor did we observe a significant correlation between discrimination performance during practice and individual congruency effects. Rather, our results suggest that blind participants were better able to discount spatial information from tactile distractors than sighted individuals, and, accordingly, take into account only the task-relevant spatial information of the target stimulus. Improved selective attention in early blind individuals has previously been reported in auditory, tactile, and auditory-tactile tasks (Collignon, Renier, Bruyer, Tranduy, & Veraart, 2006; Hötting, Rösler, & Röder, 2004; Röder et al., 1999). In the tactile domain, this performance difference has been associated with modulations of early ERP components evident in early blind but not in sighted individuals (Forster, Eardley, & Eimer, 2007). These previous studies employed deviant detection paradigms, in which participants were presented with a rapid stream of stimuli; they reacted towards stimuli matching a specified criterion while withholding responses to non-matching stimuli. As the response criterion usually comprised stimulus location, improved performance in these tasks has often been discussed in reference to altered spatial perception in blind individuals (Hötting et al., 2004). In the present study, the advantage of congenitally blind participants relates to improved selection of a target based on a non-spatial feature; given that targets could occur at any of four locations across the workspace, the improvement of selective attention in congenitally blind individuals was not tied to space. Thus, congenitally blind individuals appear to have advantages over sighted individuals on both spatial and non-spatial processing levels of tactile processing.

### 4.3. Comparison of sighted participants’ susceptibility to task instruction with previous tactile localization studies

A previous study tested sighted participants with a similar tactile congruency task as that of the present study and reported that the congruency effect always depended on the external spatial location of tactile stimuli, independent of task instructions when participants responded with the feet (Gallace et al., 2008). With the same kind of foot responses, the present study demonstrated that the direction of congruency effects is instruction-dependent. This difference may best be explained by the influence of vision on tactile localization in sighted participants: participants had their eyes open in the study of Gallace and colleagues, but were blindfolded in the present study. Non-informative vision (Newport, Rabb, & Jackson, 2002) as well as online visual information about the crossed posture of the own as well as of artificial rubber hands (Azañón & Soto-Faraco, 2007; Cadieux & Shore, 2013; Pavani, Spence, & Driver, 2000) seem to evoke an emphasis of the external reference frame. Online visual information about the current hand posture may, thus, have led to the prevalence of the externally coded congruency effect in the study by Gallace and colleagues. In contrast, blindfolding in the present study may have reduced the vision-induced bias towards external space, and, thus, allowed expression of task instruction-induced biases.

### 4.4. Lack of evidence for an effect of movement context on congruency effects

Based on previous findings with other tactile localization paradigms that have manipulated the degree of movement during tactile localization tasks (Azañón et al., 2015; Heed, Möller, et al., 2015; Hermosillo et al., 2011; Mueller & Fiehler, 2014a, 2014b; Pritchett et al., 2012), we had expected that frequent posture change would emphasize the weighting of an external reference frame in both sighted and blind participants. The reason for the lack of a movement context effect in the present study may be related to the employed distractor interference paradigm used by us. The need to employ selective attention to suppress task-irrelevant distractor stimuli might have led, as a side effect, to a discounting of task-irrelevant factors, such as movement context, in our task. Furthermore, the movements in our task differed from those of previous studies in that they only changed hand orientation, but did not modulate hand position. In contrast, other studies have always required either an arm movement that displaced the hands, or eye and head movements that changed the relative spatial location of touch to the eyes and head.

### 4.5. Summary and conclusion

In sum, we report that both sighted and congenitally blind individuals can flexibly adapt the weighting of anatomical and external information during the encoding of touch, evident in the dependence of tactile congruency effects on task context. Our results demonstrate, first, that congenitally blind individuals do not rely on only a single, anatomical reference frame but, instead, flexibly integrate anatomical and external spatial information, indicating that the use of external spatial information in touch does not ultimately depend on the availability of vision during development. Second, absence of vision during development appears to improve the ability to select tactile features independent of stimulus location, as demonstrated by reduced distractor interference in congenitally blind participants.

## FUNDING AND ACKNOWLEDGEMENTS

Data and analysis scripts are available on https://osf.io/ykqhd/. We thank Maie Stein, Julia Kozhanova, and Isabelle Kupke for their help with data acquisition, Helene Gudi and Nicola Kaczmarek for help with participant recruitment, Janina Brandes and Phyllis Mania for fruitful discussions, and Russell Lenth and Henrik Singmann for help with mixed models for data analysis. This work was supported through the German Research Foundation’s (DFG) Research Collaborative SFB 936 (project B1), and Emmy Noether grant He 6368/1-1, by which TH is funded. SB is supported by a research fellowship from the German Research Foundation (DFG), BA 5600/1-1.

